## Supplementary material for "One-trial social learning of vervet monkey alarm calling"

^*^ Correspondence:

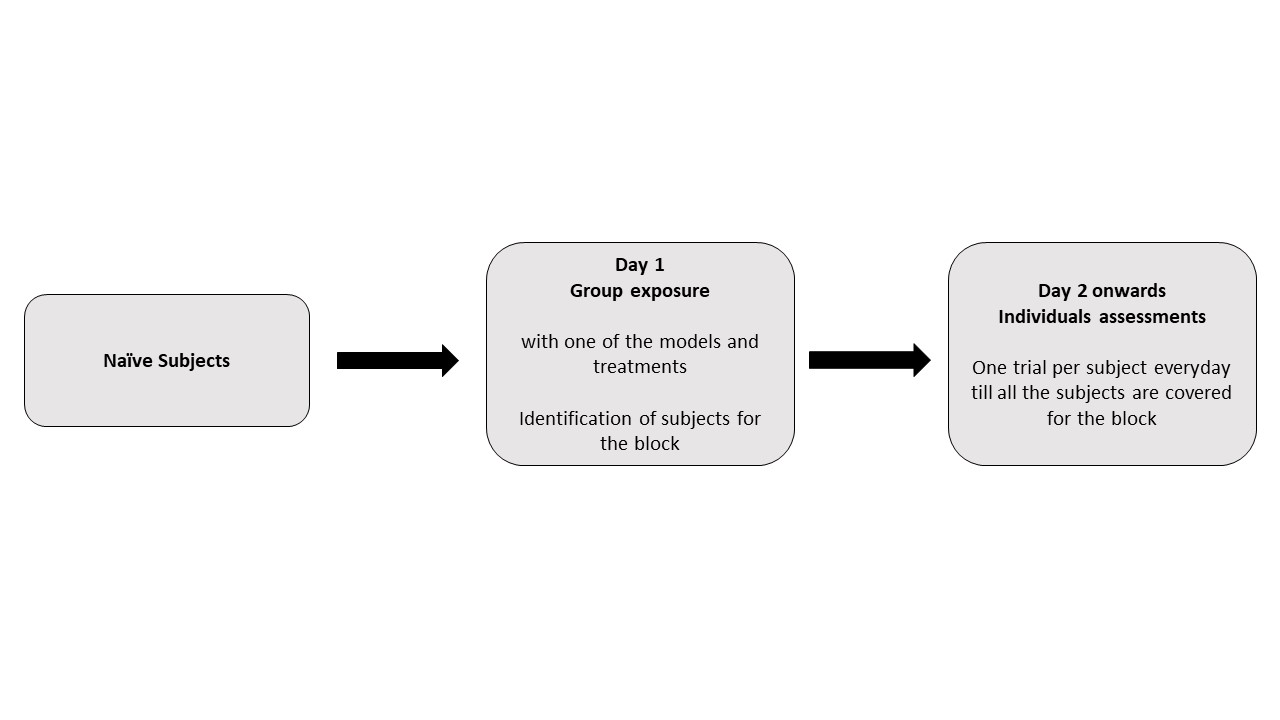

Figure S1: Design and timeline of the experimental blocks

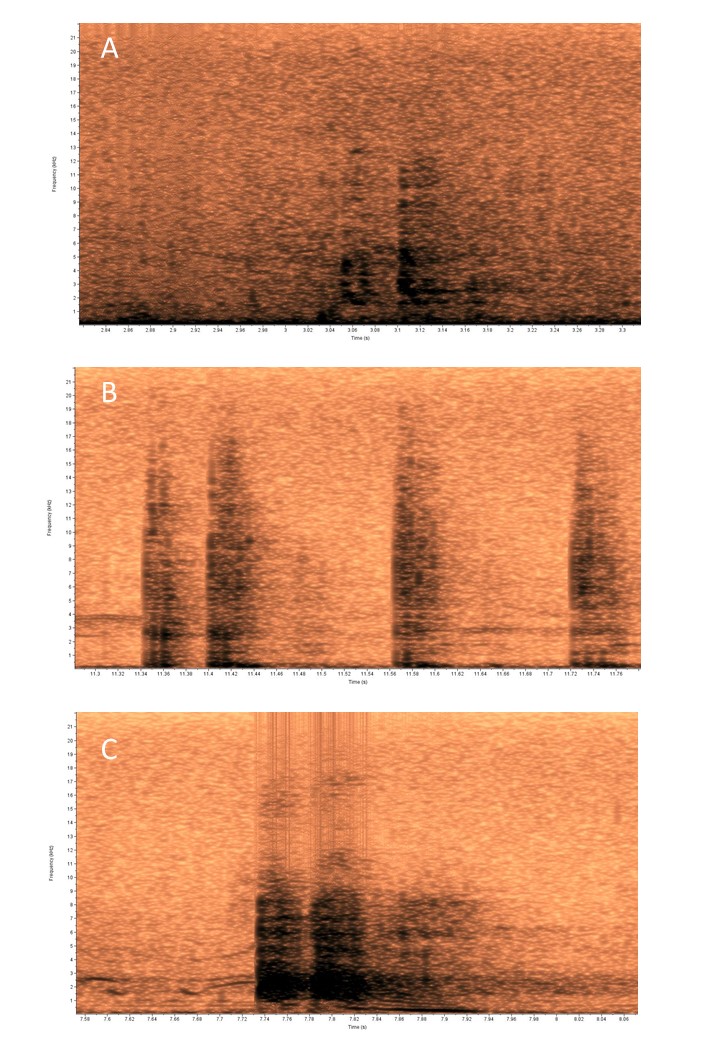

Figure S2: Spectrograms (Hamming window at 1024 DFT and 93.8% overlap) of different types of alarm calls elicited by the subjects (A) Eagle alarms (B) Snake alarms (C) Leopard alarms as described in the previous studies.

Figure S3: Mean (± SE) predator inspection of the subjects for the Grunt playback and leopard alarm playback (Alarm) treatment.

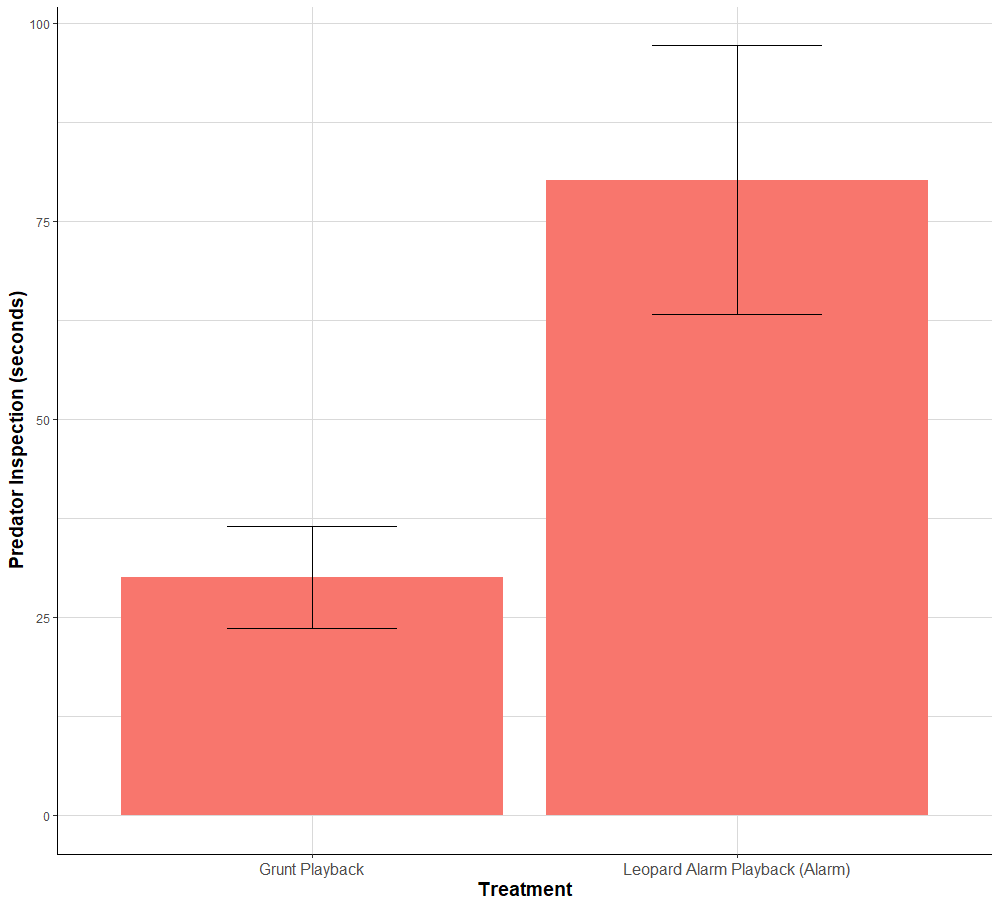

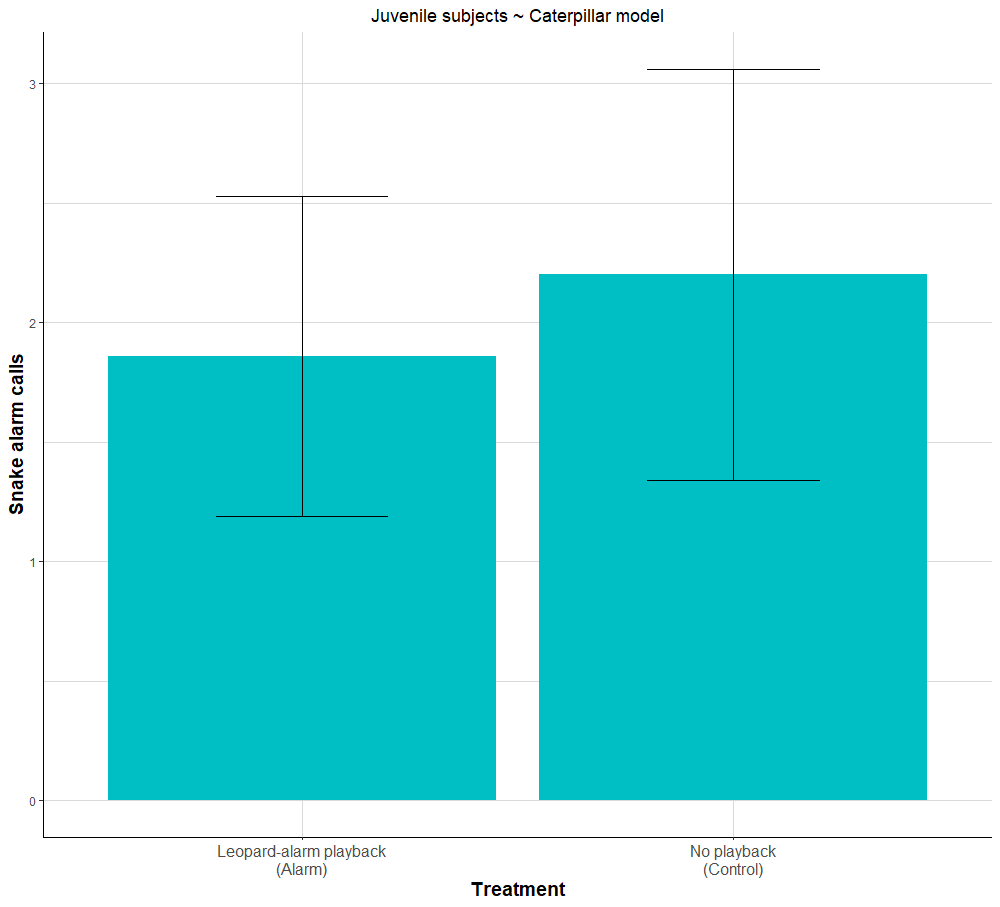

Figure S4: Mean (± SE) of snake alarms for the juvenile subjects exposed to caterpillar model

Table S1: List of acoustic parameters extracted from the alarm calls. Shaded parameters were used for the LDA classification

| **Acoustic parameter (unit)** | **Group Means from LDA Model (Scaled Data)** | | |
| --- | --- | --- | --- |
|  | Eagle | Leopard | Snake |
| Number of elements | -0.0350025 | -0.3522358 | 1.24903709 |
| Low Frequency (Hz) | 0.0736336 | -0.3854407 | 1.3558483 |
| High Frequency (Hz) | NA | NA | NA |
| Delta Time (s) | -0.0970311 | 1.89210804 | - 0.3294874 |
| Peak Frequency (Hz) | NA | NA | NA |
| 1st Quartile Frequency (Hz) | -0.908022 | -0.755146 | 1.005444 |
| 3rd Quartile Frequency (Hz) | -1.298510 | -1.112019 | 1.152288 |
| Delta Frequency (Hz) | -0.4686237 | 0.1796256 | 0.5083690 |
| Average Entropy (bits) | -1.5461260 | -1.2993005 | 0.9196337 |
| Center Frequency (Hz) | -1.130832 | -0.958378 | 1.084521 |
| Max Entropy (bits) | -1.458680 | -1.389708 | 0.633118 |
| Average Power (dB) | -0.187820 | 1.892933 | -0.514142 |
| Average Amplitude (U) | -0.07130897 | 0.34673157 | -0.07008640 |
| Max Amplitude (U) | NA | NA | NA |
| Min Amplitude (U) | 0.3045821 | -1.5637610 | 0.3033251 |
| Min Entropy (bits) | -0.9943584 | -1.1550906 | 1.0071179 |
| Energy (dB) | NA | NA | NA |
| Peak Amplitude (U) | NA | NA | NA |
| Peak Power (dB) | NA | NA | NA |
| Max Frequency (Hz) | -0.7533995 | -0.6241039 | 1.0150978 |
| Max Power (dB) | NA | NA | NA |
| Peak Frequency Contour Max Frequency (Hz) | -1.0853231 | -0.8660863 | -0.8660863 |
| Peak Frequency Contour Average Slope (Hz/ms) | -0.038994987 | -0.008381332 | -0.008381332 |
